## Supplementary Information for "An open-source closed-loop Virtual Reality system to investigate social interactions and collective behavior in fish"

### A low-cost closed-loop Virtual Reality system to investigate social interactions and collective behavior in fish
### – Supplementary information –

Stéphane Sanchez[1], Ramón Escobedo[1, 2, 3, 4], Renaud Bastien[1, 2], Boris Lenseigne[5], Audrey Denis[2], Mathieu Moreau[2], Maud Combe[2], Andrew D. Straw[6, 7], Clément Sire[4], Guy Theraulaz[2*]

[1]Institut de Recherche en Informatique de Toulouse (IRIT), Université Toulouse Capitole, Toulouse, France.
[2]Centre de Recherches sur la Cognition Animale, Centre de Biologie Intégrative, CNRS, Université de Toulouse III – Paul Sabatier, 31062, Toulouse, France.
[3]Laboratoire de Physique Théorique, CNRS, Université de Toulouse III – Paul Sabatier, 31062, Toulouse, France.
[4]Departamento de Matemáticas, Universidad Carlos III de Madrid, 28911, Leganés, Madrid, Spain.
[5] Izital BV, Delft, The Netherlands.
[6]Institute of Biology I, Faculty of Biology, Albert-Ludwigs-Universität Freiburg, Freiburg, Germany.
[7]Bernstein Center Freiburg, Albert-Ludwigs-Universität Freiburg, Freiburg, Germany.

#### 1 Experimental setup diagram and parts

The experimental setup frame is made of modular aluminum strut profiles (S1 Fig). The frame dimensions (width × depth × height) are 1 m × 1 m × 2 m. The fish tank top is fixed with an aluminum plate at mid-height of the frame. The depth camera is fixed (at about 50 cm above perpendicularly to the middle of the tank) with an articulated camera arm. IR filters are applied to the depth camera lenses. Eight IR lamps illuminate the tank from underneath. Each lamp is made from an IR LED with a 100 W LED heat sink and mounted on an articulated camera arm. A high-resolution, low-latency LED video projector is mounted on the frame, pointed at a mirror under the tank that reflects the projected virtual 3D scene on the tank. To prevent perturbations from external sources during experiments, three of the upper sides and the top of the frame are covered with black acrylic plates. The last side is closed with a blackout curtain that allows easy access to the fish tank and depth camera. S1 Table lists all materials and equipment used in the experimental setup.

#### 2 Software

Closed-loop interactions between real fish and virtual ones require three software:

- Acquisition and 3D tracking software that gets frames from the Intel RealSense D435 camera and tracks fish positions from them:
  https://doi.org/10.6084/m9.figshare.30188836.v1
- Trajectory simulator that simulates virtual fish behaviors, with or without interactions with real fish:
  https://doi.org/10.6084/m9.figshare.30188842.v1

- Rendering software that displays virtual fish according to the simulated positions and real fish position (to perform anamorphosis rendering):
https://doi.org/10.6084/m9.figshare.30188845.v1

The rendering software needs calibration routines and application:
https://doi.org/10.6084/m9.figshare.30188839.v1

### 3 Perimeter of a rhodonea.

In polar coordinates, the equation of a rhodonea can be written as $r(t) = R\cos(mt)$, where $m = n/d$. The arc length $ds$ is given by

$$\begin{aligned} ds &= \sqrt{r^2 + \left(\frac{dr}{dt}\right)^2}\, dt = \sqrt{R^2\cos^2(mt) + R^2m^2\sin^2(mt)}, \\ &= R\sqrt{1 + (m^2-1)\sin^2(mt)}. \end{aligned}$$

The perimeter is thus

$$P = \int_0^{d\pi} \sqrt{r^2 + \left(\frac{dr}{dt}\right)^2}\, dt = R\int_0^{d\pi} \sqrt{1 + (m^2-1)\sin^2(mt)}\, dt.$$

Changing variables $k = mt$, we have $dt = \frac{1}{m}dk$, and when $t$ goes from $0 \to d\pi$, then $k$ goes from $0 \to md\pi = n\pi$. Thus,

$$P = R\frac{d}{n}\int_0^{n\pi} \sqrt{1 + \left(\frac{n^2}{d^2} - 1\right)\sin^2 k}\, dk.$$

As the integrand has period $\pi$ in $k$ and $n$ is odd, $\int_0^{n\pi} dk = n\int_0^{\pi} dk = 2n\int_0^{\pi/2} dk$, so

$$P = 2Rd\int_0^{\pi/2} \sqrt{1 + \left(\frac{n^2}{d^2} - 1\right)\sin^2 k}\, dk,$$

which can be calculated as an elliptic integral of the second kind,

$$E(a) = \int_0^{\pi/2} \sqrt{1 - a^2\sin^2 k}\, dk.$$

In Rose 1, $n = 3$ and $d = 5$, and for Rose 2, $n = 3$ and $d = 1$, so the perimeters are

$$P_1 = 10R \times E\left(\frac{4}{5}\right) \quad \text{and} \quad P_2 = 2R \times E\left(\frac{2\sqrt{2}}{3}\right),$$

where, for $P_2$, we have used the transformation $E\big(\sqrt{-a^2}\big) = \sqrt{1+a^2}E\big(\sqrt{a/(1+a^2)}\big)$. For $R = 19\,\text{cm}$, the values are $P_1 \approx 126.92\,\text{cm}$ and $P_2 \approx 242.5\,\text{cm}$.

### 4 Supplementary Videos

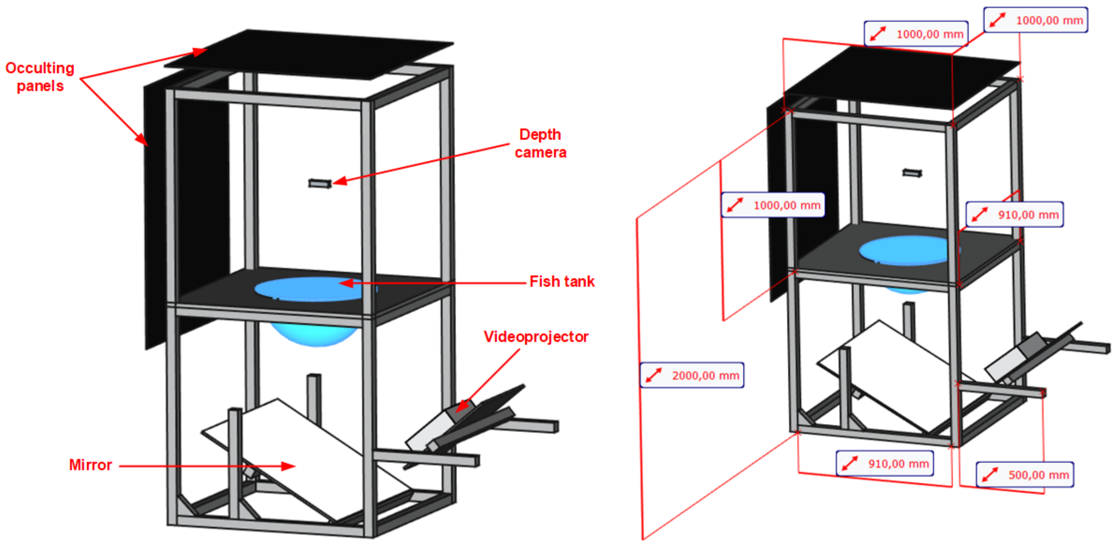


**S1 Fig. Experimental setup.** Descriptive diagram (left) and dimensions (right).

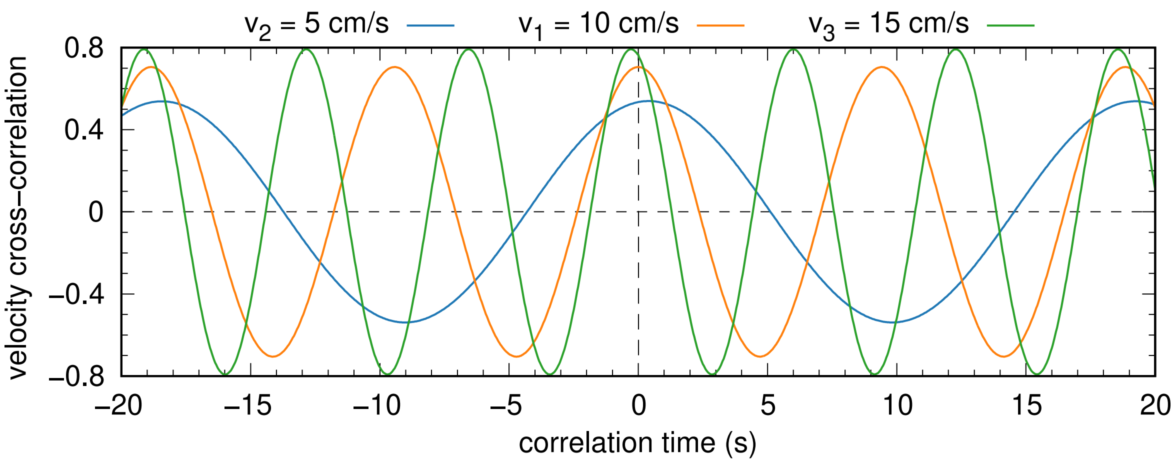


**S2 Fig. Cross-correlation of the velocity vector of the real fish with that of the virtual fish at different virtual fish speeds.** Virtual fish conditions correspond to C1, C2, and C3, with $v_1 = 10$ (orange), $v_2 = 5$ (blue), and $v_3 = 15\,\mathrm{cm/s}$ (green). The trajectories of the virtual fish are circles of radius $R = 15\,\mathrm{cm}$ at depth $z = 5\,\mathrm{cm}$ corresponding to a distance to the wall of $r_\mathrm{w} = 5.4\,\mathrm{cm}$. Maximum correlation is reached at $t_1 = 0\,\mathrm{s} + nT_1$, $t_2 = 0.4\,\mathrm{s} + nT_2$, and $t_3 = -0.267\,\mathrm{s} + nT_3$, where $T_1 = 3\pi\,\mathrm{s}$, $T_2 = 6\pi\,\mathrm{s}$, and $T_3 = 2\pi\,\mathrm{s}$, for $n = 0, \pm1, \pm2, \ldots$

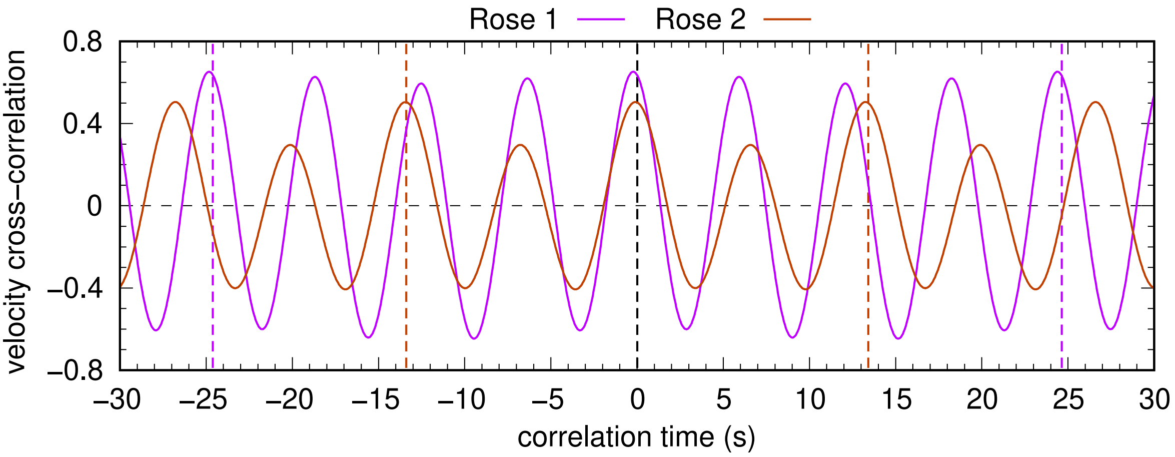


**S3 Fig. Cross-correlation of the velocity vectors of the real and virtual fish for the two rhodonea trajectories.** In both cases, the swimming speed is v = 10 cm/s, the minimum distance to the wall is $r_w = 1.4$ cm, and the swimming depth is $z = 5$ cm (corresponding to a maximum radius of $R = 19$ cm). The maximum correlation for Rose 1 (purple) is reached at $t_1 = -0.23\,s + nT_1$, with $T_1 = 24.6$ s, and for Rose 2 (brown) at $t_2 = -0.1\,s + nT_2$, with $T_2 = 13.4$ s, for $n = 0, \pm1, \pm2, \ldots$. Vertical dashed lines indicate the respective period lengths $T_1$ and $T_2$.

### 5 Supplementary Tables

| **Item Description** | **Quantity** |
|---|---|
| Aluminum profile 45 × 45 4 slots c - 2 m | 4 |
| Aluminum profile 45 × 45 4 slots 10 MM - 1 m / 91 cm | 16 |
| Aluminum profile 45 × 45 4 slots 10 MM - 0.5 m | 4 |
| Long mounting bracket for 45 × 45 profiles + screw + cover | 30 |
| Protective cap for aluminum profiles 45 × 45 10 MM slots | 12 |
| Angle bracket for 45 × 45 profiles | 8 |
| Center screw for 10 MM slot profile | 20 |
| Post-assembly fastening nuts with retaining function for 10 mm slot profiles - M8 Thread | 100 |
| Dome-head fastening screw - Thread M8x16 - Hex socket head | 100 |
| Washer 18 × 8 × 1.5 | 100 |
| Blackout acrylic panel 1 × 1 × 1.2 m | 3 |
| Blackout acrylic panel 1 × 1 × 1 m | 1 |
| Blackout curtain | 1 |
| Acrylic bowl - 50 cm diameter - 15 l contenance | 1 |
| Depth camera Realsense D435 | 1 |
| Computer - at least Intel i7 13th gen processor, 32 Gb RAM, Nvidia RTX2080 graphic card | 1 |
| 4k LED, low latency videoprojector | 1 |
| Mirror | 1 |
| IR LED lamp with 100W LED heat sink | 8 |
| Articulated camera arm | 9 |

**S1 Table. Main frame materials of the Virtual Reality setup.**

| Observables | **Conditions** (Mean ± std) | | | **Hellinger distance** | | |
|---|---|---|---|---|---|---|
| | C1 | C2 | C3 | C1\|C2 | C1\|C3 | C2\|C3 |
| Distance between fish (cm) | 9.6 ± 8.9 | 7.7 ± 7.5 | 9.7 ± 8.2 | 0.088 | 0.076 | 0.12 |
| Speed of the real fish (cm/s) | 9.4 ± 3.6 | 6.3 ± 3.1 | 11.7 ± 4.5 | **0.333** | **0.262** | **0.484** |
| Depth of the real fish (cm) | 4.4 ± 1.3 | 4.4 ± 1.1 | 5.1 ± 1.8 | 0.074 | 0.186 | 0.164 |

**S2 Table. Hellinger distance between the probability distribution function (PDF) of the 3 observables used to quantify the impact of virtual fish's swimming speed on real fish's behavior.** We list the distances obtained for the three conditions compared pairwise. The real fish maintained the same swimming distance and depth relative to the virtual fish across the three swimming speeds tested (i.e., no significant differences were found between the PDFs). However, the swimming speeds of the real fish were significantly different between the three conditions C1, C2, and C3 (i.e., when the virtual fish moved at 5, 10, and 15 cm/s, respectively). Values of Hellinger distance are shown in bold font when $H > 0.2$ (high dissimilarity of the PDFs).

| Observables | **Conditions** (Mean ± std) | | | **Hellinger distance** | | |
|---|---|---|---|---|---|---|
| | C1 | C8 | C9 | C1\|C8 | C1\|C9 | C8\|C9 |
| Distance between fish (cm) | 9.6 ± 8.9 | 7.5 ± 7.6 | 5.4 ± 5.0 | 0.118 | **0.221** | 0.139 |
| Speed of the real fish (cm/s) | 9.4 ± 3.6 | 9.2 ± 3.5 | 9.4 ± 3.1 | 0.067 | 0.113 | 0.111 |
| Depth of the real fish (cm) | 4.4 ± 1.3 | 3.9 ± 1.1 | 7.6 ± 1.4 | **0.224** | **0.7** | **0.774** |

**S3 Table. Hellinger distance between the probability distribution function (PDF) of the 3 observables used to quantify the impact of virtual fish's swimming depth on real fish's behavior.** We list the distances obtained for the three conditions compared pairwise. The swimming distance of the real fish to the virtual fish was significantly smaller when the virtual fish was at 8 cm depth. There was also a significant difference in the swimming depth of the real fish between the three conditions C1, C8, and C9. Values of Hellinger distance are shown in bold font when $H > 0.2$ (high dissimilarity of the PDFs).

| Observables | **Conditions** (Mean ± std) | | | **Hellinger distance** | | |
|---|---|---|---|---|---|---|
| | C1 | C6 | C7 | C1\|C6 | C1\|C7 | C6\|C7 |
| Distance between fish (cm) | 9.6 ± 8.9 | 6.3 ± 5.7 | 10.1 ± 9.3 | 0.192 | 0.078 | **0.208** |
| Speed of the real fish (cm/s) | 9.4 ± 3.6 | 10.1 ± 3.4 | 8.6 ± 3.6 | 0.084 | 0.08 | 0.154 |
| Depth of the real fish (cm) | 4.4 ± 1.3 | 4.4 ± 1.0 | 4.4 ± 1.3 | 0.105 | 0.101 | 0.124 |

**S4 Table. Hellinger distance between the probability distribution function (PDF) of the 3 observables used to quantify the impact of the virtual fish's distance to the wall on the real fish's behavior.** We list the distances obtained for the three conditions compared pairwise. The swimming speed and depth of the real fish were not influenced by the virtual fish's proximity to the tank wall. Only when the virtual fish was swimming close to the wall at 2.4 cm in condition C7, the real fish maintained a higher distance in comparison with condition C6, in which the distance to the wall of the virtual fish was much larger (10.4 cm). Values of Hellinger distance are shown in bold font when $H > 0.2$ (high dissimilarity of the PDFs).
